# Predicting potential current distribution of *Lycorma delicatula* (Hemiptera: Fulgoridae) using MaxEnt model in South Korea

**DOI:** 10.1101/557421

**Authors:** Hyeban Namgung, Min-Jung Kim, Sunghoon Baek, Joon-Ho Lee, Hyojoong Kim

## Abstract

*Lycorma delicatula* (Hemiptera: Fulgoridae) is invasive insect in Korea which causes plant damages by sucking and sooty molds. *Lycorma delicatula* was first detected in South Korea in 2004, where its introduction and spreading possibly were affected by human activity-related factors. Here, we used MaxEnt to describe current distribution of *L. delicatula* in Korea and tried to find out the impact of human influences for distribution. We used 143 sites of occurrence data, 19 bioclimatic variables, duration of temperature below −11°C, average daily minimum temperature in January, cumulative thermal unit variable, the distribution of grape orchard variable and human footprint to create models. These models were estimated by two sets of 24 candidates with feature combinations and regularization multipliers. In addition, these two sets were created as models with and without footprint for how human influence affect to distribution. Model selection for optimal model was performed by selecting a model with a lowest sum of each rank in small sample-size corrected Akaike’s information criterion and difference between training and test AUC. Model of LQ10 parameter combinations was selected as optimal models for both model sets. Consequently, both of distribution maps from these models showed similar patterns of presence probability for *L. delicatula*. Both models expected that low altitude regions were relatively more suitable than mountain areas in Korea. Footprint might be limited for the distribution and *L. delicatula* might already occupy most of available habitats. Human-related factors might contribute to spread of *L. delicatula* to uninfected areas.

## Introduction

Spotted lanternfly, *Lycorma delicatula* (Hemiptera: Fulgoridae), is an invasive species of which origin is the southern China and the countries in the subtropical zones of Southeast Asia [1]. In Korea, *L. delicatula* was first detected in Cheonan in 2004 [2], and then expanded across South Korea for more than a decade [3]. Since its first detection, the agricultural area, mostly grapevine yards, damaged by *L. delicatula* increased rapidly from one ha in 2006, seven ha in 2007, 91 ha in 2008, 2,946 ha in 2009, to 8,378 ha in 2010 [4, 5]. According to Park et al. [3], *L. delicatula* might disperse more frequently in the western region and its long-distance dispersal could be possible beyond the mountain range. These rapid spread might be caused by human-related factors such as vehicles which mainly could transfer a host plant or other material with its egg masses [6].

A few studies have been conducted for determining potential habitats and habitat suitability of *L. delicatula*. Jung et al. [7] estimated the potential habitats of *L. delicatula* in Korea by using CLIMEX, which is a mechanistic modelling method based on physiological traits and constraints [7]. The CLIMEX requires the biological parameters related with the target insects such as optimal temperature, lower developmental threshold, lethal temperature, optimal humidity and so on [8]. Nevertheless, information on the biological parameters for *L. delicatula* was limited, and thus estimated potential habitats were not exactly matched with the current distribution of *L. delicatula* in Korea. A correlative method, which relates occurrence data of a species to environmental data statistically [9], seems to be better than deterministic methods (e.g., CLIMEX) because biological information for *L. delicatula* is limited and human related factors cannot be applied to deterministic methods due to difficulty of parameterization of them. As one of the correlative methods, MaxEnt could be applied to describe current distribution of *L. delicatula* in Korea because it needs only presence data and has high predictive accuracy [10, 11], although a few issues with MaxEnt modeling should be considered to increase prediction accuracy (e.g., sampling bias of occurrence data, types of feature, model complexity, criteria of model selection, model evaluation method and etc.) [12-15].

This study aims to select best combinations of parameters for application of presence data of *L. delicatula* and its surrounding environmental and human related variables in MaxEnt, to select variables affecting the current distribution, to find out the effects of anthropogenic factor, and to visualize current presence probability of *L. delicatula* in Korea.

## Material and methods

### Collection and preparation of presence data of *L. delicatula*

Total 143 presence data points of *L. delicatula* were collected from the published papers [6, 16-19] (15 points), the report of National Institute of Ecology (NIE) [20] (83 points), and observation of this study (45 points). For developing distribution maps of *L. delicatula*, data were divided into two data sets; one for training data for model calibration, and the other for test data for model validation or evaluation [9]. For training data, 83 data points of the NIE report were selected because these data were collected for whole Korean territories with a consistent sampling criteria during one year, in 2015. The other 60 points were used for test data. Both training and test data sets were analyzed to determine spatial patterns using ArcGIS 10.1 [21] with the average nearest neighbor test, and all data were used in model development because both data show a random distribution. Data with random distribution are needed to avoid overestimating problem, which is caused by clustered occurrence data, in species distribution models [22]

### Environmental variables related to *L. delicatula*

Monthly temperature and precipitation data from 1981 to 2010 (30 years) were downloaded from the web site of the Korea Meteorological Administration (KMA). These weather data were collected from 73 meteorological stations operated nationwide by KMA, and then were interpolated by Inverse Distance Weighting (IDW) method for estimating temperature and precipitation with a grid size of 1 km. Nineteen bioclimatic variables [23] were created in DIVA-GIS 7.5 [24] using these data. These 19 variables were transformed to ASCII file format using SDMs tool [25] in ArcGIS 10.1.

To consider all variables related with occurrence of *L. delicatula*, published papers on its ecology were reviewed (Table 1). Among them, two overwintering related variables, development related variable and main host (i.e., grape) distribution of *L. delicatula*, were used as these variables expected to be directly related with occurrence of *L. delicatula* in Korea. There were multiple studies [6, 26-28] that overwintering egg mortality of *L. delicatula* was affected by number of days with minimum temperature below −11°C and average daily minimum temperature in January. Thus, these variables (i.e., under_-11_Jan and min_tmp_jan) were created with the same climatic data and interpolation method, and then used for 19 bioclimatic variables. Cumulative thermal unit variable (i.e., Degree day) for development of *L. delicatula* in locations of 73 meteorological stations of KMA was calculated and mapped by using average daily maximum and minimum temperatures of 30 years (1981-2010) and 11.13°C lower development threshold from Park’s paper [6]. Even although *L. delicatula* has diverse host plants [16], its adults showed high preference and fitness at grapes [16, 29]. Thus, distribution of grape orchard variable (i.e., Grape) was created from 1,916 dimensions of viticulture by regions (www.agrix.go.kr) with ordinary krigging method in ArcGIS 10.1.

**Table 1.**
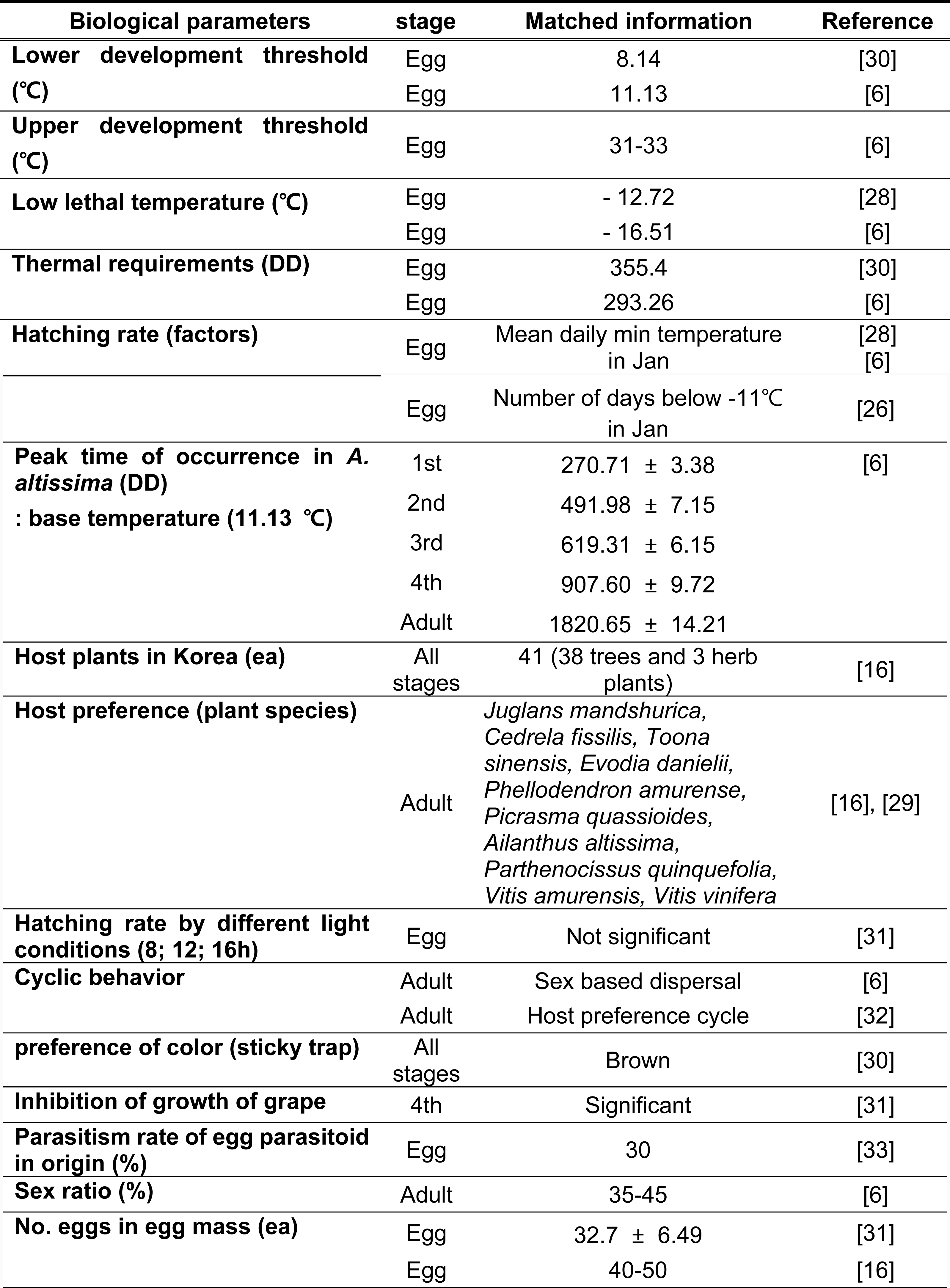
Biological information of *L. delicatula* in published papers.

Human foot print variable (i.e., footprint) [34] was also downloaded (http://sedac.ciesin.columbia.edu/) because distribution of *L. delicatula* might be affected by human activities [6].

Total 24 environmental variables (i.e., 19 bioclimatic variables, under_-11_Jan, min_tmp_jan, Degree day, Grape and footprint) were created to develop distribution model of *L. delicatula* In Korea.

### Selection of environmental variables

Multi-collinearity test was conducted to eliminate correlated variables by Pearson’s coefficient ‘*r*’ [35]. If multiple variables were correlated (| *r* | > 0.8), only one variable was selected based on biological relevance with *L. delicatula* ecology. From this process, 11 variables (bio03, bio05, bio11, bio12, bio13, bio15, bio17, under_-11_Jan, Degree day, grape and footprint) were selected among 24 variables.

### Modelling procedure

As a default setting, MaxEnt offers six features (i.e., an expanded set of transformations of the original covariates [36] types, L (linear), Q (quadratic), P (product), T (threshold), H (hinge), C (categorical), which are automatically selected by ‘Auto features’ depending on the sample size of training data [37]. In addition, MaxEnt creates a distribution model, using regularization multiplier (default value = 1), which mitigates model complexity or overfitting, to make general interpretation [36]. Nevertheless, MaxEnt does not always create the best model by a given default parameter setting [38]. Therefore, to select the best model we adjusted the parameters setting and developed 24 candidate models with different feature combinations and regularization multipliers by using four feature combinations (LQ, LQP, LQH, LQPH) and six regularization parameters (1, 2, 5, 10, 15, 20). For comparison, another set of 24 models trained by 10 variables except for footprint was also created with previously noted feature combinations and regularization parameters to estimate habitat suitability excluded possibility of propagation.

For selecting optimal parameter combination, small sample-size corrected Akaike’s information criterion (i.e., AIC_c_) [39] and area under the receiver operation characteristic curve (i.e., AUC) were used to compare candidate models. Because high training AUC (AUC calculated by training data) might be result of overfitting model, difference between training and test AUC (AUC calculated by test data) were choose for model selection criteria. Therefore, AIC_c_ and difference between training and test AUC (AUC_diff_) were calculated in ENMTools [40] and MaxEnt. To consider both AUC_diff_ and AIC_c_ [12] for model selection, sum of each rank in AUC_diff_ and AIC_c_ from the lowest value was used because smaller values of AIC_c_ and AUC_diff_ represent a better model. If sum of rank of candidate models is equal, a model with a smaller AIC_c_ score was selected. From these processes, optimal models were re-built and importance of each variables evaluated with jackknife test and 10-fold cross-validation. Each of two models were built with its own selected variables to describe current occurrence probability of *L. delicatula* in Korea. The maps of two models were visualized in ArcGIS 10.1.

## Results

### Selection of best parameter combinations in both models for MaxEnt application

LQ10 (i.e., combination of feature types L and Q and regularization multiplier 10), LQH10 (i.e., combination of feature types L, Q, and H and regularization multiplier 10), and LQPH5 (i.e., combination of feature types L, Q, P, and H and regularization multiplier 5) were selected by having the lowest value of summing both ranks in AUC_diff_ and AIC_c_ for the model without footprint (Fig 1. (A)). However, both LQ10 and LQH10 had same AIC_c_ values, and these values of both parameter combinations were lower than one of LQPH5. Thus, LQ10 and LQH10 were considered as the best models without footprint for *L. delicatula.* LQ10 and LQH10 were also determined in the model with footprint (Fig 1. (B)). In both model selections, LQ10 and LQH10 created exactly same model. Therefore, LQ10 parameter combinations of each model was selected to build the distribution model for *L. delicatula*.

**Fig 1.**
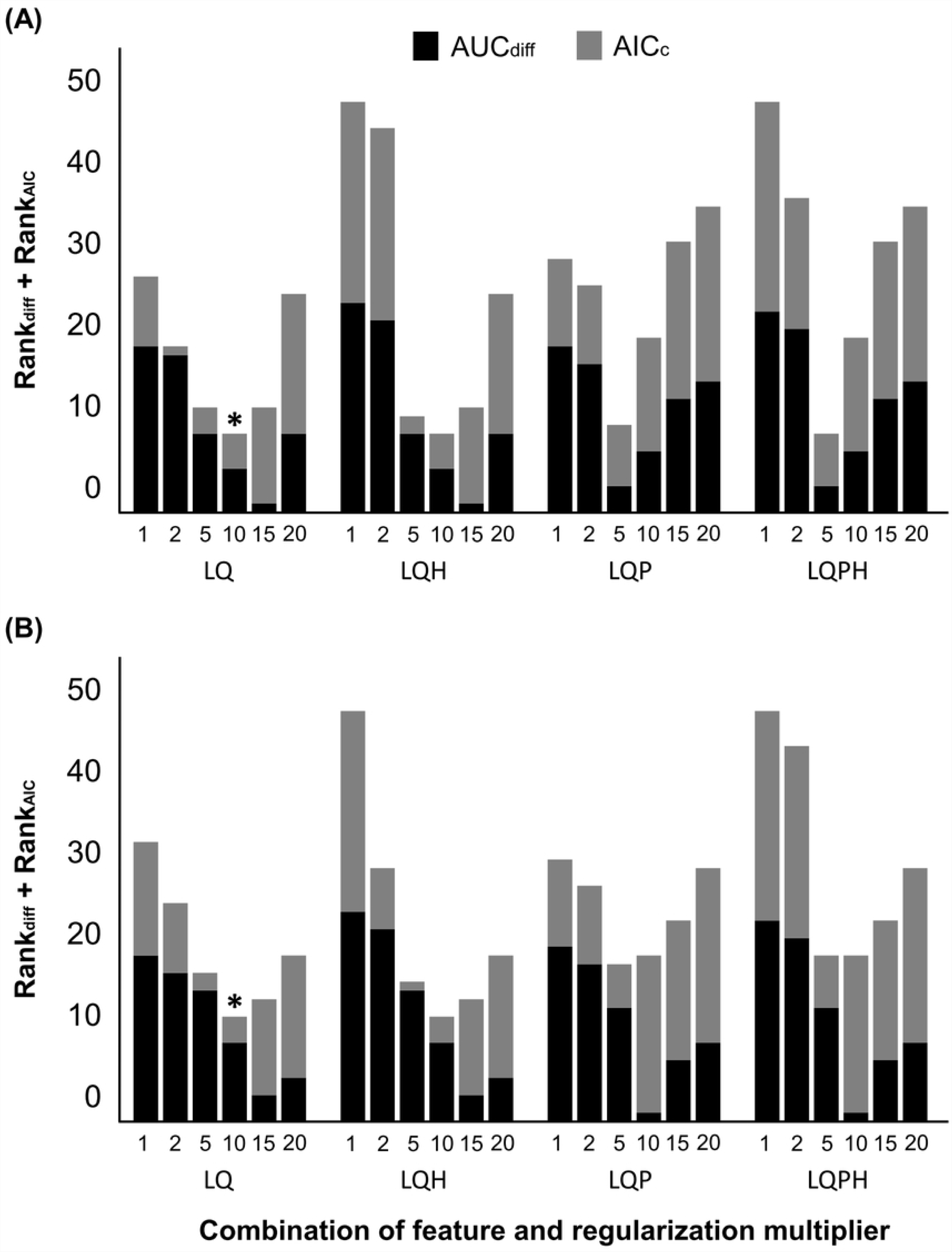
Sum of AUC_diff_ and AIC_c_ ranks for 24 candidate models. Sum of AUC_diff_ and AIC_c_ ranks of all candidate models (A) without footprint and (B) with footprint. Each AUC_diff_ and AIC_c_ were ranked in order of ascending power from values calculated in MaxEnt and ENMTools. Black and grey bar represent rank of AUC_diff_ and AIC_c_ respectively. Asterisk (*) represents final selected models.

### Evaluation of developed models

Two MaxEnt executions (without footprint and with footprint) with LQ10 parameter combination created models using five variables in each run (Table 2). The average AUC score calculated by 10-fold cross-validation with training data and 10 variables for the model without footprint was 0.733 ± 0.064 (Table 2), indicating reasonable performance (AUC score > 0.7; Peterson at al., 2011). When this model was evaluated with test data, the AUC score was 0.747, representing reliable performance. Among five variables used in modelling, Bio 05 and Degree day were estimated as important variables in distribution model for *L. delicatula* by occupying more than 90% contributions to determine the distribution model. This result was also obtained in jackknife test (Fig 2. (A) and Table 3).

**Table 2.** Summary of description and performance of each two models.

| Model | Model description | Model performance |
| --- | --- | --- |

| (LQ 10) | No. input variables | Used variables | No. parameters | Training AUC | Average AUC (mean±SD) | Test AUC |
| --- | --- | --- | --- | --- | --- | --- |
| **Without footprint** | 11 | Bio 05<br>Bio 13<br>Bio 15<br>Degree day<br>Grape | 5 | 0.755 | 0.733 ± 0.064 | 0.747 |
| **With footprint** | 10 | Bio 05<br>Bio 13<br>Degree day<br>Footprint<br>Grape | 5 | 0.789 | 0.769 ± 0.045 | 0.773 |

Training AUC and test AUC were calculated by training and test data of occurrence information respectively. Average AUC was averaged AUC value of 10 test bins in result of 10-folds cross-validation.

**Table 3.** Averaged percent contribution and permutation importance of environmental variables for each two models.

| Variables | Without footprint |  | With footprint |  |
| --- | --- | --- | --- | --- |
|  | Percent contribution | Permutation importance | Percent contribution | Permutation importance |
| **Footprint** | - | - | 51.4 | 27 |
| **Bio 05** | 91.3 | 75.2 | 40.8 | 53.3 |
| **Bio 15** | 4 | 1.9 | - | - |
| **Degree day** | 2 | 16.5 | 2.4 | 15.4 |
| **Grape** | 1.7 | 2.8 | 1.3 | 2.9 |
| **Bio 13** | 1 | 3.3 | 0.1 | 1 |

**Fig 2.**
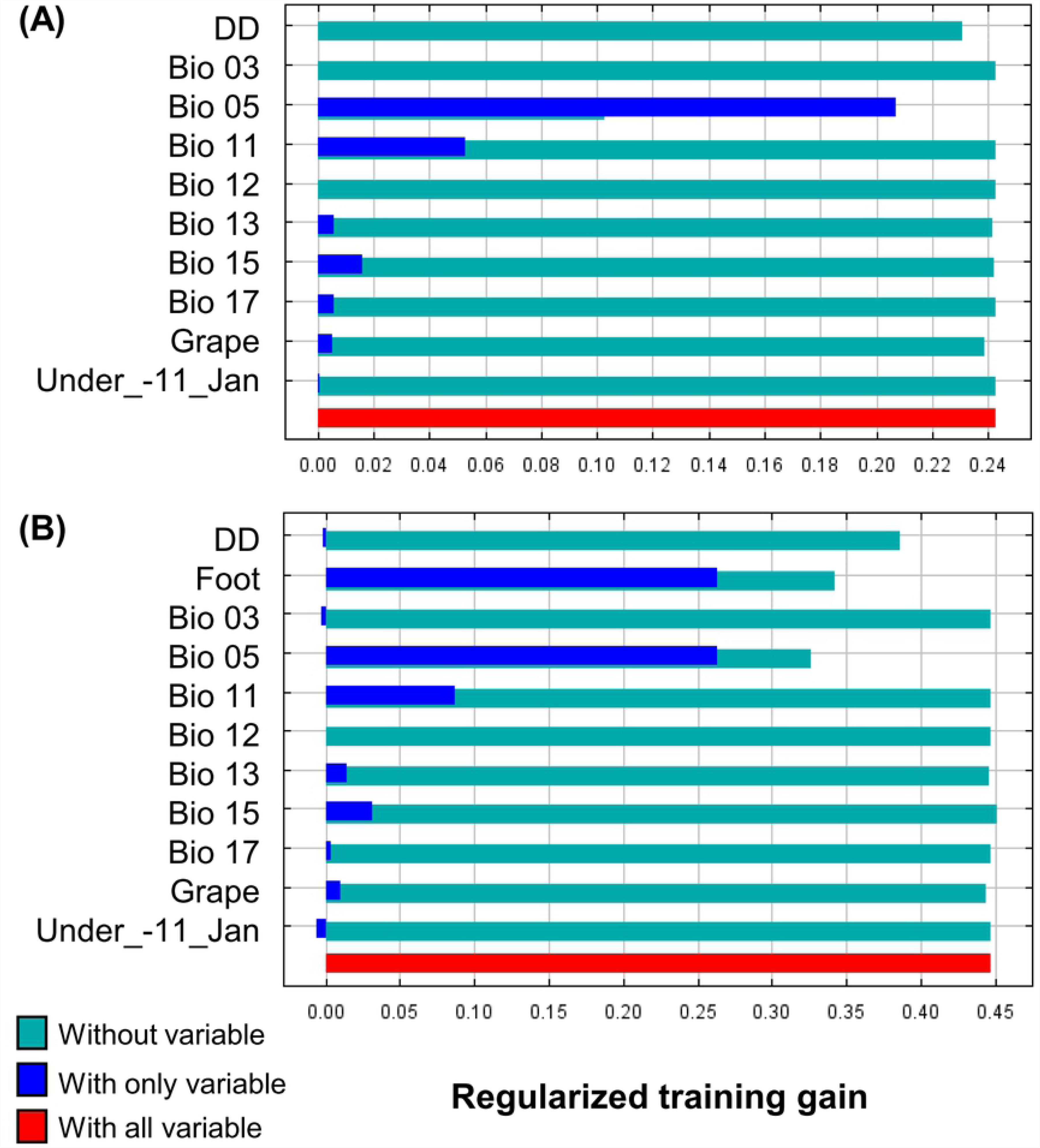
Results of Jackknife test for relative importance of used variables in each final model. Relative importance of (A) ten variables used in model without footprint and (B) 11 variables used in model with footprint. Jackknife tests were executed with 10-fold cross-validation, results were averaged values of each run.

The model with footprint performed better than the model without footprint, showing that the average AUC score was 0.769 ± 0.045 and test AUC score was 0.773 (Table 2). This model was also created with five parameters from five variables regularized by LQ 10 combination. The footprint and Bio 05 were important variables predicting distribution of *L. delicatula* (Fig 2 and Table 3). The response curve of footprint and Degree day showed that probability of presence was increased linearly (Fig 3. (A) and (C)). Probability of presence was exponentially increased according to the increase of Bio 05 (Fig 3. (B)).

**Fig 3.**
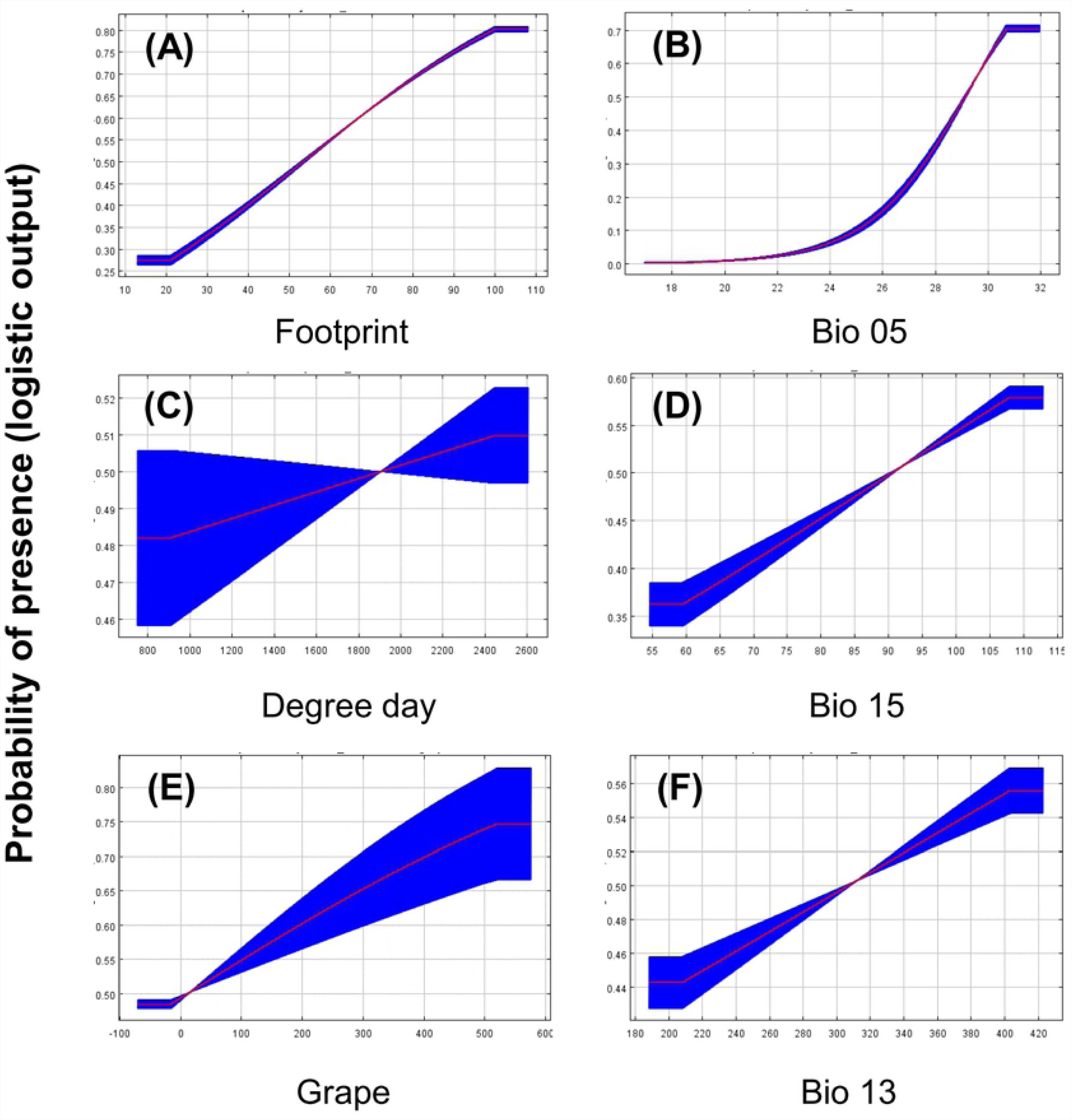
Probability of presence of *L. delicatula* according to each environmental variable. (A) Human foot print; (B) Max temperature of warmest month; (C) Cumulative thermal unit (base temperature: 11.13°C); (D) Precipitation seasonality (coefficient of variation); (E) Distribution of grape orchard; (F) Precipitation of wettest month

### Current presence probability maps of *L. delicatula* in Korea

Both distribution maps including and excluding footprint variable well descripted current presence possibility of *L. delicatula* in Korea, showing similar patterns of presence probability (Fig 4). Both models expected that mountain areas were potentially unsuitable, whereas low altitude regions were relatively suitable in Korea (Fig 4). However, the model with footprint more specifically estimated regions of higher probability of presence rather than that without footprint (Fig 4).

**Fig 4.**
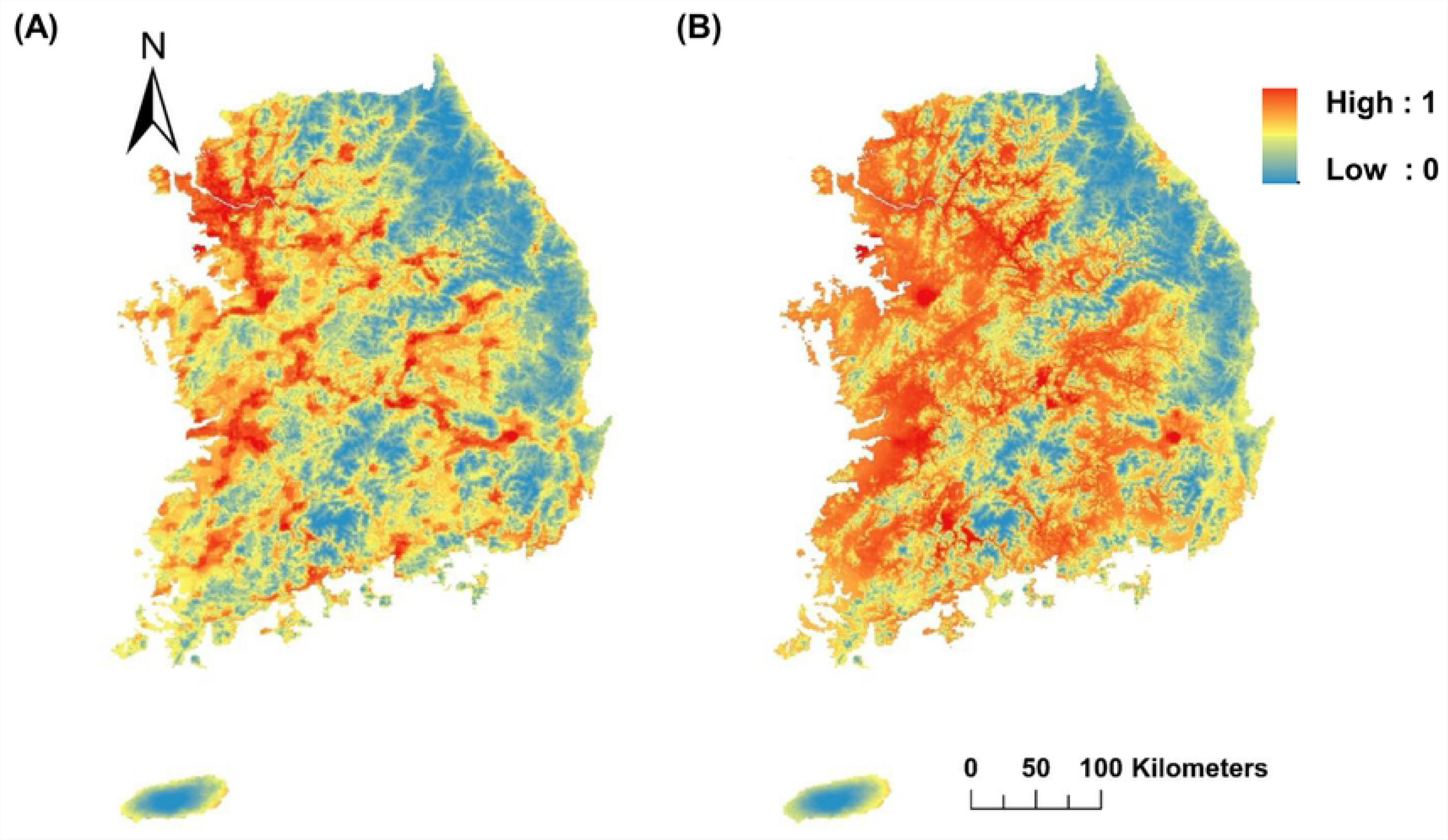
Potential distribution map of *L. delicatula* estimated by two models in Korea. Probability of presence (MaxEnt’s logistic output) predicted by (A) model with footprint and (B) without footprint.

## Discussion

Both models without and with footprint predicted well current presence possibility of *L. delicatula* in Korea. Moreover, there was a strong correlation (r = 0.916) between two models. This indicates that the effects of human-related factors might be limited for the current distribution of *L. delicatula,* which might already occupy most of available habitats in Korea. However, the model with footprint showed higher prediction ability than that without footprint by considering AUC values. From our results, we speculate that human-related factors might contribute to spread of *L. delicatula* to uninfected areas rather than directly affection for habitat suitability. Spear et al. [42] found strong relationship between human population density and richness of alien species, proposing human density as good predictor determining population size of alien species. This human-mediated propagule pressure could assist successful establishment of invasive species by supplying population above Allee threshold into new areas [43]. As for *L. delicatula*, contribution of human factors for its dispersal could be significant because its eggs were frequently found in packing, construction and agricultural materials [6, 31]. Recently, *L. delicatula* was also found in Berks county of Pennsylvania, USA [44, 45]. This introduction and spreading were also suspected by causing human-related factors such as packing materials and vehicles [44, 45]. Moreover, there was no record of *L. delicatula* found in the DMZ (Demilitarized Zone) area [46, 47] even though physical distance was not far away from the observed areas. Therefore, human influence could be an important factor in determining distribution of *L. delicatula*, especially in the early stage of invasion.

In both models with and without footprint, Bio 05 (i.e., maximum temperature during the warmest month) was the most important environmental factor to contribute the distribution of *L. delicatula*. This might be related to the origin of *L. delicatula* which is South China and Southeast Asia [48]. Because *L. delicatula* is poikilothermic, its development is increased as temperature increases up to its upper developmental threshold [49-51]. The mean maximum temperature during the warmest month in Korea was generally lower than the upper developmental threshold of *L. delicatula* [6, 52].

Degree day, another environmental factor made in this study, also contributes in distribution modeling of *L. delicatula*. In response curve of Degree day variable, 50% presence possibility of *L. delicatula* was determined around 1,900 DD similar to the accumulated degree-days (i.e., 1,821) of peak occurrence of *L. delicatula* adults in fields [6]. One-tailed binomial tests [53] were applied to training and test data using 1,821 DD, the Degree day variable well distinguished presence and pseudo-absence of *L. delicatula* significantly (*p* < 0.05) in both training and test data. Therefore, degree-days would be suitable variable to predict potential habitat of *L. delicatula*.

The other variables (i.e., Bio 13, Bio 15, and grape) used in modeling contribute a small amount and discrimination ability of these variables for presence or absence of *L. delicatula* was very low. The five variables related to winter temperature (Bio 06, Bio 09, Bio 10, under_-11_Jan, and min_tmp_jan), which are supposed to determine the hatching rate of *L. delicatula*, were not selected for the distribution model of *L. delicatula*. Although these variables were proposed as important variable determining annual population size in many papers [6, 26-28], they did not explain distribution of *L. delicatula* in this study. This suggests that low lethal temperature (i.e., around −12.7 °C to −16.5 °C) is over than average winter temperature in Korea, like Bio 05. As an example, there was no case that January mean temperature was less than −11 °C in Seoul, one of coldest areas in Korea, during 38 years (1981-2018), and quite high as −2.56 ± 1.951 °C (mean ± SD) than lower lethal temperature.

Both models in this study are closer to realized niche than fundamental one because these models were not built by deterministic method finding physiological traits of *L. delicatula*, but had correlative method with distribution and environmental variables [9]. These two models are strictly realized niche only in Korea, which include environmental variables in Korea and their unknown interaction [54]. Thus, extrapolation to other regions or predict of future distribution needs cautions. However, it could be still applicable to predict risk analysis for *L. delicatula* even in non-contaminated areas and countries having high risk of being invaded.

In conclusion, major variables related with occurrence of *L. delicatula* in Korea should be helpful for predicting its occurrence. Moreover, footprint variable might be applicable for making surveillance plan and deciding domestic quarantine stations in countries with early stages of invasion of *L. delicatula*, while remaining relevant variables with the occurrence of *L. delicatula* could be used for risk assessment in non-invaded countries

## Acknowledgements

This work was carried out with the support of “Cooperative Research Program for Agriculture Science & Technology Development (Project No. PJ01257203)” Rural Development Administration, Republic of Korea. And this work was supported by Korea Environment Industry & Technology Institute (KEITI) through Exotic Invasive Species Management Program, funded by Korea Ministry of Environment (MOE) (2018002270005).

